# Improving qBOLD based measures of oxygen extraction fraction using hyperoxia-BOLD derived measures of blood volume

**DOI:** 10.1101/2020.06.14.151134

**Authors:** Alan J Stone, Nicholas P Blockley

## Abstract

**Purpose:** Streamlined-qBOLD (sqBOLD) is a refinement of the quantitative BOLD (qBOLD) technique capable of producing non-invasive and quantitative maps of oxygen extraction fraction (OEF) in a clinically feasible scan time. However, sqBOLD measurements of OEF have been reported as being systematically lower than expected in healthy brain. Since the qBOLD framework infers OEF from the ratio of the reversible transverse relaxation rate (R_2_′) and deoxygenated blood volume (DBV), this underestimation has been attributed the overestimation of DBV. Therefore, this study proposes the use of an independent measure of DBV using hyperoxia-BOLD and investigates whether this results in improved estimates of OEF.

**Methods:** Monte Carlo simulations were used to simulate the qBOLD and hyperoxia-BOLD signals and to compare the systematic and noise related errors of sqBOLD and the new hyperoxia-qBOLD (hqBOLD) technique. Experimentally, sqBOLD and hqBOLD measurements were performed and compared with TRUST (T_2_ relaxation under spin tagging) based oximetry in the sagittal sinus.

**Results:** Simulations showed a large improvement in the uncertainty of DBV measurements leading to a much improved dynamic range for OEF measurements with hqBOLD. In a group of ten healthy volunteers, hqBOLD produced measurements of OEF in cortical grey matter (OEF_hqBOLD_ = 38.1 ± 10.1 %) that were not significantly different to TRUST oximetry measures (OEF_TRUST_ = 40.4 ± 7.7 %), whilst sqBOLD derived measures (OEF_sqBOLD_ = 16.1 ± 3.1 %) were found to be significantly different.

**Conclusion:** The simulations and experiments in this study demonstrate that an independent measure of DBV provides improved estimates of OEF.

## Introduction

Oxygen extraction fraction (OEF) is an important indicator of the metabolic function of brain tissue that describes the fraction of oxygen removed from arterial blood to serve oxidative metabolism. The quantitative BOLD (qBOLD) technique provides the capability to non-invasively and quantitatively map OEF on a regional level^1,2^. Streamlined-qBOLD (sqBOLD) is a refinement of this approach that seeks to simplify the acquisition and analysis of data to provide a more clinically relevant approach^3^.

Measurements of OEF using sqBOLD are lower than expected when compared with literature values (OEF_sqBOLD_ ∼ 20%, OEF_literature_ ∼ 30 - 40%)^3^. Since OEF is derived from the ratio of the irreversible transverse relaxation rate (R_2_′) and the deoxygenated blood volume (DBV), it follows that the underestimation of OEF could be due to underestimation of R_2_′ or overestimation of DBV. The sqBOLD technique utilises an asymmetric spin echo (ASE) acquisition, with which R_2_′ has been measured to be in the range of 2.6–3.6 s^−1^ in grey matter (GM)^3–5^. This is consistent with previous ASE measurements^6^ of 3.5 s^−1^ and with different acquisitions such as gradient echo sampling of spin echo (GESSE)^2^ measuring 2.9 s^−1^. DBV has been measured by sqBOLD to be in the range 3.6–6.7 % in GM^3–5^. This is consistent with other ASE based measurements^6^ of 4.3 %, but not with previous measurements using GESSE^2^ of 1.8 % or hyperoxia-BOLD^7^ of 2.2 %, suggesting that DBV overestimation might be responsible for OEF underestimation in ASE based qBOLD. This hypothesis is given further weight by simulations of ASE based qBOLD showing that diffusion, which is not incorporated in the qBOLD model, causes DBV to be overestimated^8^. There is therefore great potential to improve the accuracy of sqBOLD by improving the accuracy of DBV measurements.

Multiparametric qBOLD (mq-BOLD) introduced the concept of acquiring an independent measurement of blood volume which is combined with a measurement of R_2_′ acquired from separate maps^9,10^ of T_2_ and T_2_*. Here the blood volume is measured using the dynamic susceptibility contrast (DSC) technique^11^ which requires an injection of a Gadolinium-based contrast agent. However, the DBV in the qBOLD model explicitly refers to the proportion of the blood volume that contains deoxyhaemoglobin. Whereas DSC is sensitive to all of the vascular compartments that the contrast agent passes through and hence is a measure of total cerebral blood volume (CBVt). Therefore, for sqBOLD this DSC measure of CBV would likely drive OEF estimates even lower.

Hyperoxia-BOLD provides an alternative to DSC which is targeted specifically at measuring venous CBV (CBVv). It has been shown that the fractional change in the BOLD signal in response to the administration of oxygen is specific to CBVv and can be scaled to provide quantitative estimates using a heuristic model^7^. The hyperoxia-BOLD and qBOLD techniques rely on the same BOLD contrast. The former uses oxygen as a contrast agent by manipulating the concentration of deoxyhaemoglobin in the deoxygenated blood vessels, whilst the latter relies on the additional signal attenuation due to deoxyhaemoglobin at short ASE refocussing offset times. Therefore, theoretically the CBVv from hyperoxia-BOLD is equivalent to DBV from qBOLD and henceforth we will refer to both as DBV.

The aim of this study is to investigate the use of hyperoxia-BOLD to improve the accuracy of OEF measurements. Firstly, detailed simulations of the proposed technique were performed using a Monte Carlo based approach^8^. The potential for systematic error due to physiological variability and the effect of noise were both examined and compared with ASE based qBOLD. Secondly, experimental measurements were performed to compare the existing sqBOLD approach with the new hyperoxia-qBOLD (hqBOLD) technique. Furthermore, comparison is made of both techniques with whole brain oximetry measurements from the TRUST (T_2_ relaxation under spin tagging) technique^12^.

## Methods

### Simulations

Simulations of the qBOLD and hyperoxia-BOLD signal were performed using a previously reported approach^8^. A brief summary of this approach follows. Monte Carlo simulations of the random walk of protons around deoxygenated blood vessels modelled as infinite cylinders were performed using a standard approach^13^ for a range of different vessel radii and nominal oxygenation at 3 T. The resulting phase accrual was saved for each proton in 2 ms intervals for a total of 120 ms. The ASE signal is then generated by summing the phase accrued prior to the refocussing pulse and subtracting the phase accrued following the refocussing pulse. Whilst the gradient echo (GRE) signal is produced by summing the phase accrued up to the required TE. These data can be scaled to the target oxygenation^14^ prior to summation of the individual proton’s phases to simulate the extravascular signal. This extravascular signal is then used to estimate a radius dependent shape function enabling the signal to be scaled to the target blood volume fraction^15^. To account for the range of vessel radii encountered *in vivo*, the extravascular signal from multiple vessel radii can be combined as the product of single radii simulations^15,16^. A discrete model of the cerebral vasculature was used to select these radii^17^. Finally, the intravascular signal is simulated using an analytical model of the blood signal^18,19^.

In this study, simulations of the ASE qBOLD and GRE BOLD signals were performed to match the experimental parameters described in later sections. Due to the limitations of the existing Monte Carlo data, the simulated spin echo displacement times (*τ*) of the ASE measurements were required to be a multiple of 4 ms and the TE of the BOLD measurements a multiple of 2 ms. Since the experimental values did not meet these criteria a wider range of *τ* and TE values were simulated and linearly interpolated to the required values. The hyperoxia-BOLD experiment was simulated by estimating the oxygen carrying capacity (CaO_2_) of arterial blood during normoxia and hyperoxia,

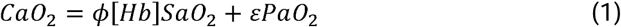

where the constant *φ* is the oxygen carrying capacity of haemoglobin (1.34 mlO_2_ g^−1^), [Hb] is the haemoglobin concentration (typical value 15 gHb dl^−1^), *ε* is the solubility coefficient of oxygen in plasma (0.0031 mlO_2_ dl^−1^ mmHg^−1^) and the arterial oxygen saturation (SaO_2_) was estimated using the Severinghaus equation^20^. The amount of oxygen extracted for metabolism (CmetO_2_) was assumed to be constant for normoxia and hyperoxia, enabling the venous oxygen carrying capacity (CvO_2_) to be estimated.

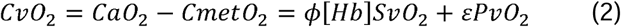

The venous partial pressure of oxygen (PvO_2_) was calculated assuming that the plasma oxygen component was non-negligible and used to calculate the venous oxygen saturation (SvO_2_) via the Severinghaus equation. Finally, the capillary oxygen saturation (ScO_2_) was calculated as a weighted sum of the SaO_2_ and SvO_2_ (ScO_2_ = *κ* SaO2 + (1-*κ*) SvO_2_), with a weighting factor *κ*=0.4 giving a greater weighting towards venous blood^21,22^.

The combined physical and physiological model of the MRI signal was used in two ways: (i) to investigate the presence of systematic error and (ii) to investigate the effect of noise on the parameter estimates. Systematic error was investigated in the absence of system noise and incorporated variations in CBVt, OEF, haematocrit (Hct), normoxic PaO_2_ (PaO ^norm^) and hyperoxic PaO_2_ (PaO_2_^hyper^). As in previous publications, this was achieved by randomly generating values in a predefined range (Table 1) using a uniform random number generator for all five parameters^23^. These values were then used to generate ASE and BOLD signals using the model. This process was repeated to produce 1,000 different physiological states and accompanying signals. These signals were processed to generate estimates of R_2_′ and OEF using the process described in the analysis section below. The estimation of DBV differed from the experimental method, which uses linear regression, in that single estimates of the BOLD signal at normoxia and hyperoxia were used to calculate the percentage change in the BOLD signal (*δ*BOLD) between these two conditions.

**Table 1:** Simulation parameters used to define the physiological state. The standard value was used when investigating the effect of noise and the range tested reflects the ranges used to investigate systematic error.

| Parameter | Standard Value | Range tested | Description |
| --- | --- | --- | --- |
| CBVt | 5 % | 0 – 10 % | Total cerebral blood volume |
| OEF | 40 % | 0 – 100 % | Oxygen extraction fraction |
| Hct | 34 % | 31 – 43 % | Small vessel haematocrit |
| PaO<sub>2</sub><sup>norm</sup> | 110 mmHg | 100 – 130 mmHg | Normoxic arterial partial pressure of oxygen |
| PaO<sub>2</sub><sup>hyper</sup> | 400 mmHg | 370 – 450 mmHg | Hyperoxic arterial partial pressure of oxygen |

The effect of noise was investigated by simulating signals for standard physiological values (Table 1). Gaussian random noise was then added to these signals before calculating R_2_′, DBV and OEF. This process was repeated 20,000 times. The SNR for ASE and BOLD was estimated for the whole brain from the experimental data following Method 1 of the National Electrical Manufacturers Association (NEMA) guidelines^24^. For the ASE data two consecutively acquired *τ*=0 (spin echo) images were used. Across the group, ASE SNR was measured as 52 and for BOLD the SNR was 87. To account for the averaging process inherent in the experimental linear regression analysis, multiple signals with added noise were generated for normoxia and hyperoxia and averaged before calculating *δ*BOLD. This consisted of 100 signal averages during hyperoxia and 100 during normoxia.

For the purposes of comparison, the *true value* of a parameter is required. This is problematic given the definition of DBV as the vascular volume containing deoxygenated blood since this applies to both capillaries and veins. However, the capillaries contain proportionally less deoxyhaemoglobin due to a higher oxygen saturation and generate a smaller fraction of the overall qBOLD signal i.e. the signal from capillaries is expected to increase quadratically with blood oxygenation, but the linear scaling constant is 100 times smaller than for large venous vessels^25^. Therefore, pragmatically DBV is defined as the blood volume within the venous vessels of the model. This is consistent with our previous simulations of the hyperoxia-BOLD signal^7^, but not our previous simulations of the qBOLD signal^8^. The true R_2_′ was then calculated using this definition of DBV,

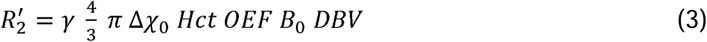

where *γ* is the proton gyromagnetic ratio, Δ*_x_*_0_ is the susceptibility difference between oxygenated and deoxygenated red blood cells (Δ*_x_*_0_ = 0.264 × 10^−6 26^), B_0_ is the main magnetic field (B_0_ = 3 T) and the known Hct from the simulations. The true OEF was an input to the simulations.

### Imaging

Ten healthy participants (aged 23 – 41; median 29; 3 female, 7 male) were scanned with local ethics committee approval using a 3T Siemens Prisma (Siemens Healthcare, Erlangen, Germany) with a 32-channel receive-only head coil. Figure 1 presents an overview of the imaging data acquired, the parameters derived from this imaging data and the main comparisons performed in this study. MRI data were acquired in the following order:

**Figure 1:**
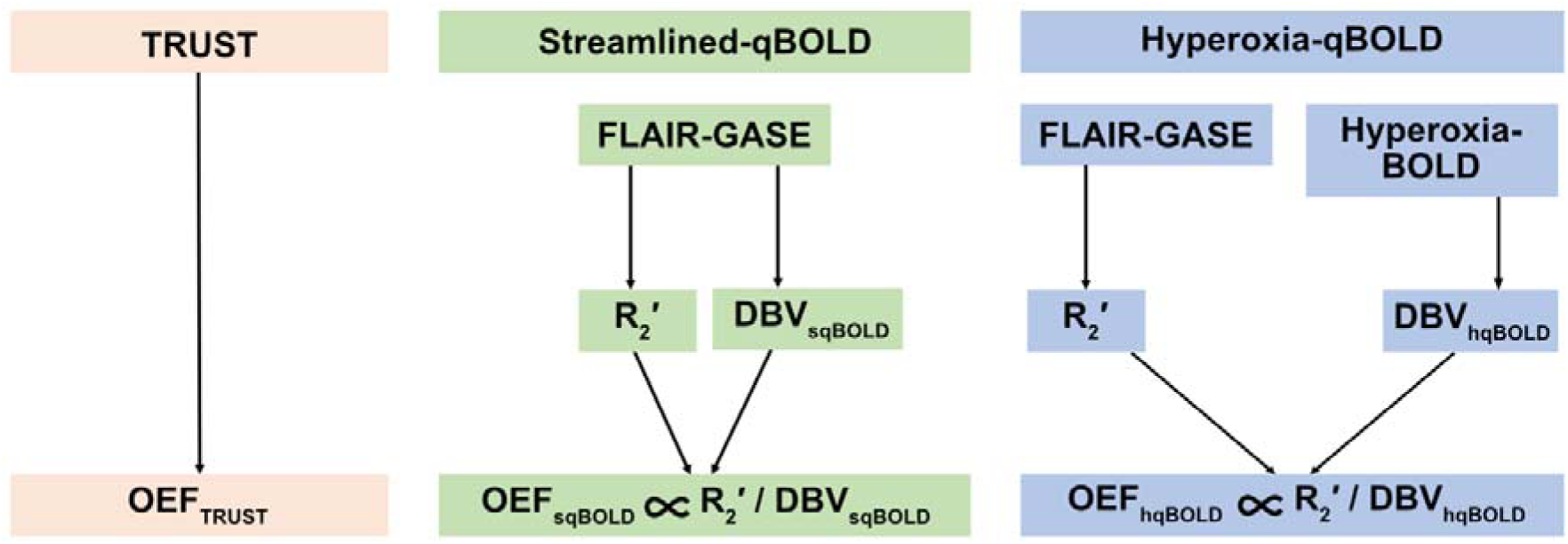
Schematic outlining the MR / physiological parameters derived from each imaging method which include the TRUST (T_2_ relaxation under spin tagging) method for measuring whole brain oxygen extraction fraction (OEF), the streamlined-qBOLD technique for mapping tissue OEF and deoxygenated blood volume (DBV) and the hyperoxia-qBOLD approach which uses an independent measurement of DBV from hyperoxia BOLD to map OEF.

TRUST oximetry measurements were made in the superior sagittal sinus (SSS) using the following parameters: FOV = 230 mm^2^, 64 x 64 matrix, TR / TE = 3 s / 7 ms, flip angle (FA) = 90*°*, GRAPPA = 3, partial Fourier = 6/8, bandwidth (BW) = 2604 Hz/px, tag-gap = 25 mm, tag-thickness = 100 mm, TI = 1050 ms. Four tag–control pairs were acquired at four different effective echo times (eTE = 0, 40, 80 and 160 ms) resulting in thirty-two acquisitions and a total scan duration of 1:53 mins.

The sqBOLD measurements were made using a FLAIR-GASE acquisition^27,28^ with the following parameters: FOV = 220 mm^2^, 96 x 96 matrix, nine 5 mm slabs (encoded into four 1.25mm slices, 100% partition oversampling), 2.5 mm gap, TR / TE = 3 s / 80 ms, FA = 90*°*, BW = 2004 Hz/px, TI_FLAIR_ = 1210 ms, τ = 0 – 66 ms in steps of 3 ms. This resulted in twenty-three τ-weighted acquisitions with a total scan duration of 9:12 mins.

The same ASE data were used for the hqBOLD measurements and complemented with hyperoxia-BOLD data to independently estimate DBV. BOLD-EPI data were acquired with a matched slice prescription to the FLAIR-GASE acquisitions (FOV = 220 mm^2^, 96 x 96 matrix, nine 5 mm slices, 2.5 mm gap, TR / TE = 1 s / 35 ms, FA = 65*°*, BW = 2004 Hz/px). A prospective end-tidal gas targeting system (RespirAct^TM^ Gen 3, Thornhill Research Inc., Toronto, Canada) was used to modulate end-tidal oxygen (PETO_2_) between normoxic and hyperoxic (baseline + 300 mmHg) conditions, whilst maintaining isocapnia (a constant carbon dioxide level). The hyperoxia paradigm lasted 10 minutes during which time 600 volumes were acquired. The respiratory paradigm consisted of three 2 minute blocks of normoxia interleaved with two 2 minute blocks of hyperoxia. The combined acquisition time was 19:12 mins.

An MPRAGE dataset (FOV = 174 mm x 192 mm x 192 mm, 116 x 128 x 128 matrix, TR / TI / TE = 1900 / 904 / 3.74 ms and FA = 8°) was acquired in each subject^29^. To aid in the registration of the FLAIR-GASE to the structural image, an additional set of whole brain GASE data without a FLAIR preparation were acquired with an increased coverage in the z direction and a set of ASE data with τ = 0 to estimate image SNR.

### Data Analysis

The following pre-processing and analysis steps were applied to the data acquired in each subject using a combination of tools from the FMRIB Software Library (FSL)^30^ and custom scripts written using MATLAB (The Mathworks, Natick, MA).

The tag and control images of the TRUST data were first motion corrected using the FMRIB linear image registration tool (MCFLIRT) (Jenkinson et al., 2002). Pairwise subtraction of the tag-control images was performed and the four repeats at each eTE averaged. Using the difference image at eTE = 0 ms, the four voxels with the highest signal in the SSS were identified to create a region of interest (ROI). Using the SSS ROI, the mean difference signal (ΔS) was extracted for each eTE. To calculate the T_2_ of blood (T_2b_), ΔS is plotted as a function of eTE and fitted to Eq. 4 to obtain the exponent C ^12^.

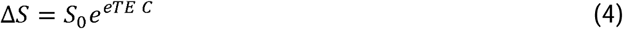

The T_2_ of blood (T_2b_) was then estimated from C using Eq. 5 and assuming the T_1_ of blood (T_1b_) to be 1624 ms.

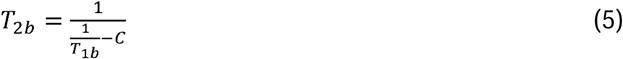

Assuming a value for haematocrit in large vessels (Hct = 0.42)^31^, T_2b_ can be converted to SvO_2_ using a calibration model^32^. Assuming arterial blood is fully saturated, an estimate of whole brain OEF is given by 1-SvO_2_.

For sqBOLD, the four 1.25 mm slices of each slab of the FLAIR-GASE data were averaged to produce nine 5 mm slices, which were then motion corrected using MCFLIRT^33^. A mask of brain tissue was created using the brain extraction tool (BET)^34^ and further analysis was restricted to voxels within this mask. The τ-series for each voxel were then fit using a linear system (A · x = B) to simultaneously estimate R_2_′, DBV and a constant term representing the underlying proton density and T_2_ decay^4^.

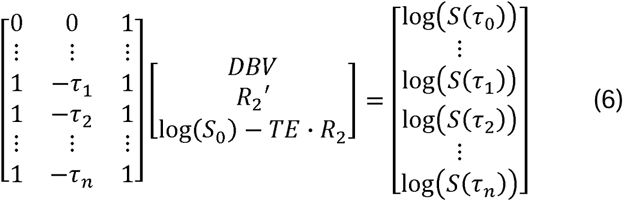

Here, S(*τ*) is the signal intensity as a function of *τ*, where *τ*_0_ represents acquisitions at *τ* = 0 ms and *τ*_1_ to *τ*_n_ are values of τ > 15 ms. In matrix A the first row describes the short τ regime relevant to τ_0_ and subsequent rows the long *τ* regime^35^ representing *τ* > 15 ms. The least squares solution was used to produce voxel-wise estimates of R_2_′ and DBV. Parameter maps of OEF were then calculated using Δ*_x_*_0_ = 0.264 × 10^−6 26^ and small vessel Hct^36^ of 0.34.

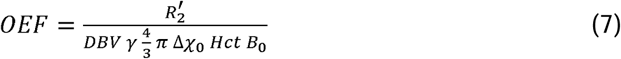

For the multi-parametric hqBOLD approach, a map of R_2_′ was calculated from the same data as for sqBOLD but only for images with *τ* > 15 ms using a log-linear least squares fit for R_2_′ and the constant term. The hyperoxia-BOLD data were then analysed in the following way. The BOLD data were motion corrected using MCFLIRT and then transformed into the FLAIR-GASE image space using FLIRT^37^. Measurements of PETO_2_ were interpolated onto the 1 sec time resolution of the BOLD data and smoothed to generate a regressor of the blood oxygenation change. This model was fitted to the hyperoxia-BOLD data on a voxel-wise basis using the least squares method and used to calculate the fractional BOLD signal change to hyperoxia (*δ*BOLD). The change in PETO_2_ during the hyperoxia challenge (ΔPETO_2_) was calculated by taking the mean of data within windows during baseline (repetitions 1 to 100) and hyperoxia (repetitions 175 to 225 and 415 to 465). These measurements were used to calculate DBV, under the assumption that ΔPETO_2_= ΔPaO_2_ (the change in PaO_2_) using Eq. 8, where the constant A = 27 ms, B = 0.2, C = 245.1 mmHg, D = 0.1 have been previously described^7^.

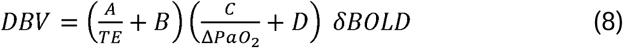

### Regional Analysis

Subject specific ROIs of cortical GM and white matter (WM) were created from the FLAIR-GASE (*τ*=0) data using FMRIB’s automated segmentation tool (FAST)^38^ by taking advantage of the T_1_-weighting caused by the FLAIR preparation. GM partial volume estimate (PVE) maps were threshold at 50% and further refined to cortical GM by transforming the MNI structural atlas^39^ into the FLAIR-GASE space via the MPRAGE using FLIRT. Frontal, insula, occipital, parietal and temporal lobes were included in the cortical mask to produce the final cortical GM ROI. WM ROIs were generated by thresholding the WM PVE maps at 100%.

### Statistical testing

A one-way ANOVA was used to test the null hypothesis of no difference between the OEF estimates made using the three different techniques. Post-hoc statistical testing was performed using the Tukey Kramer method (honest significant difference test) to test for differences between the estimates made using the different methods. A paired t-test was used to test the null hypothesis of no difference between the DBV estimates made using sqBOLD and hyperoxia-BOLD.

## Results

Table 2 displays ΔPETO_2_ and the change in end-tidal partial pressure of carbon dioxide (ΔPETCO_2_) during the hyperoxia-BOLD experiment for all participants. The mean ΔPETO_2_ was 294.7 mmHg and the mean ΔPETCO_2_ was 0.4 mmHg.

**Table 2:** Parameter estimates of deoxygenated blood volume (DBV) and oxygen extraction fraction (OEF) calculated using streamlined-qBOLD (sqBOLD), hyperoxia-qBOLD (hqBOLD) and T_2_ relaxation under spin tagging (TRUST). Median values in global grey matter are shown for each subject and presented alongside the group mean and standard deviation.

| Subject | PET<sub>O<sub>2</sub></sub> (mmHg) | ΔPET<sub>O<sub>2</sub></sub> (mmHg) | PET<sub>CO<sub>2</sub></sub> (mmHg) | ΔPET<sub>CO<sub>2</sub></sub> (mmHg) |
| --- | --- | --- | --- | --- |
| sub-01 | 122.9 | 321.5 | 34.9 | 1.0 |
| sub-02 | 108.9 | 277.8 | 41.7 | 1.2 |
| sub-03 | 109.8 | 295.6 | 43.3 | 0.5 |
| sub-04 | 109.2 | 297.0 | 41.3 | -0.2 |
| sub-05 | 104.9 | 291.0 | 39.6 | -0.1 |
| sub-06 | 109.3 | 268.2 | 39.5 | 2.3 |
| sub-07 | 112.0 | 296.9 | 36.3 | -0.5 |
| sub-08 | 128.8 | 312.6 | 35.4 | 0.5 |
| sub-09 | 128.8 | 277.7 | 37.0 | 0.6 |
| sub-10 | 109.0 | 308.7 | 42.2 | -1.1 |
| Mean | 114.4 | 294.7 | 39.1 | 0.4 |
| SD | 8.9 | 16.8 | 3.0 | 1.0 |

Figure 2 presents the results of the simulations of the sqBOLD and hqBOLD techniques. The R_2_′ estimated from the simulated qBOLD data is plotted against the true R_2_′ based on the pragmatic definition of DBV from above (Fig. 2a). A dashed line of unity is plotted and the slope of the relationship between the data is estimated as 0.99 (intercept -0.24 Hz). DBV_sqBOLD_ shows a large amount of uncertainty when compared with true DBV, which increases with true OEF (Fig. 2b). Whilst DBV_hqBOLD_ has a much lower uncertainty when OEF > 30 % (Fig. 2c). The slope of the relationship between DBV_hqBOLD_ and true DBV is 0.78 (intercept 0.05 %) for the full range of OEF values or 0.94 (intercept 0.01 %) when OEF > 30 %. OEF_sqBOLD_ does increase with true OEF, but has a very small dynamic range (Fig. 2d). Finally, OEF_hqBOLD_ increases monotonically with true OEF for OEF > 20 % with a dynamic range consistent with the full range of OEF (Fig. 2e).

**Figure 2:**
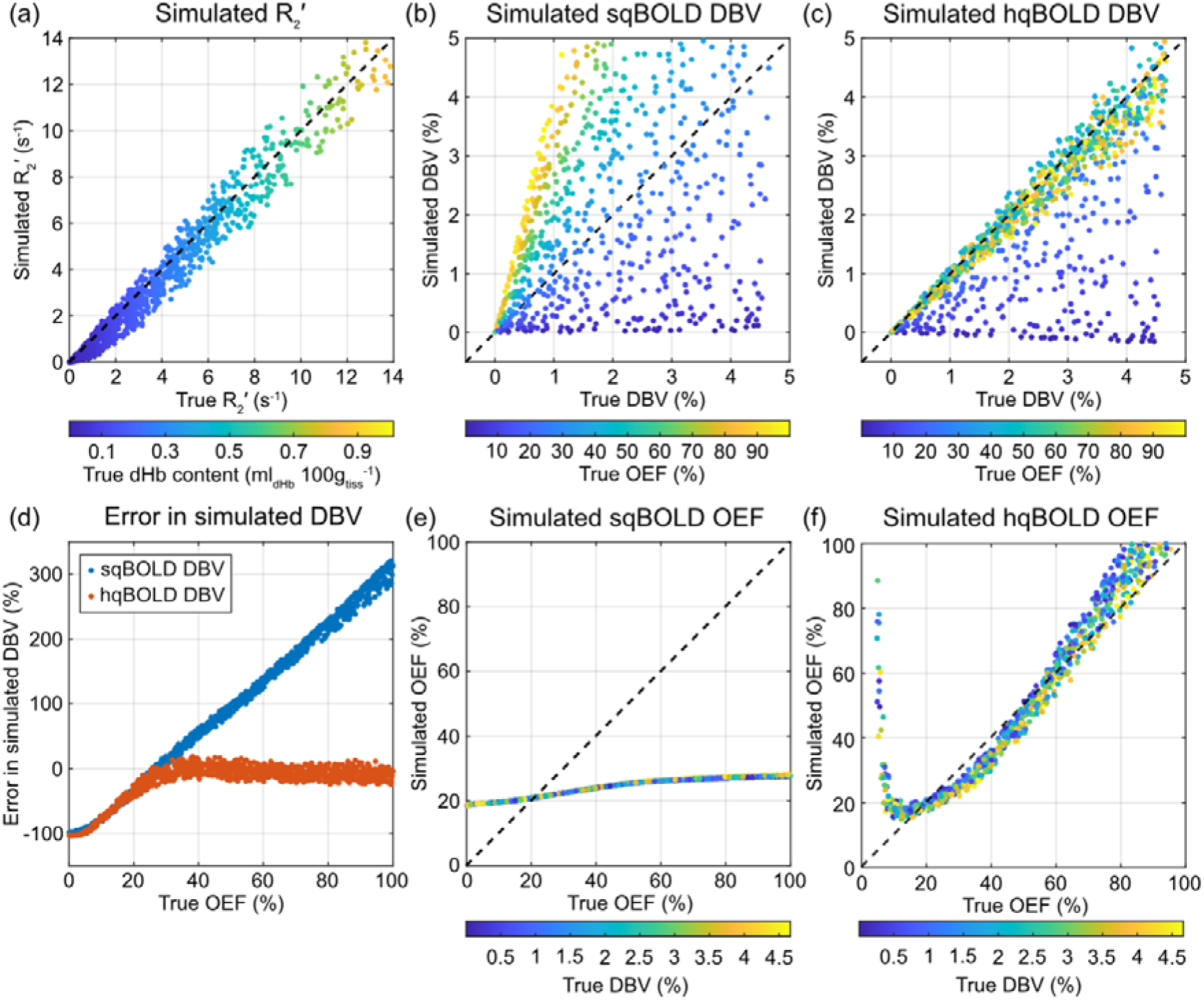
Simulations of the systematic error in the two qBOLD approaches. (a) simulated R_2_′ compared with ‘true’ R_2_′ calculated using the static dephasing qBOLD model, (b-c) simulated streamlined-qBOLD (sqBOLD) DBV and hyperoxia-BOLD (hqBOLD) DBV vs true DBV, respectively, (d) error in simulated DBV for sqBOLD and hqBOLD techniques and (e-f) simulated OEF from sqBOLD and hqBOLD, respectively.

Figure 3 presents simulations of the effect of system noise on measurements made using sqBOLD and hqBOLD. The true R_2_′ was estimated to be 2.8 s^−1^, whilst the median value simulated was 2.3 s^−1^ (Fig. 3a). DBV_sqBOLD_ has an interquartile range (IQR) of 3.2 % (Fig. 3b), compared with the DBV_hqBOLD_ IQR of 0.27 % (Fig. 3c). The main difference in the sqBOLD and hqBOLD estimates of OEF is in their propensity for outliers, with the former having an excess kurtosis value of 17577.5 (Fig. 3d) and the latter a value of 0.1 (Fig. 3e).

**Figure 3:**
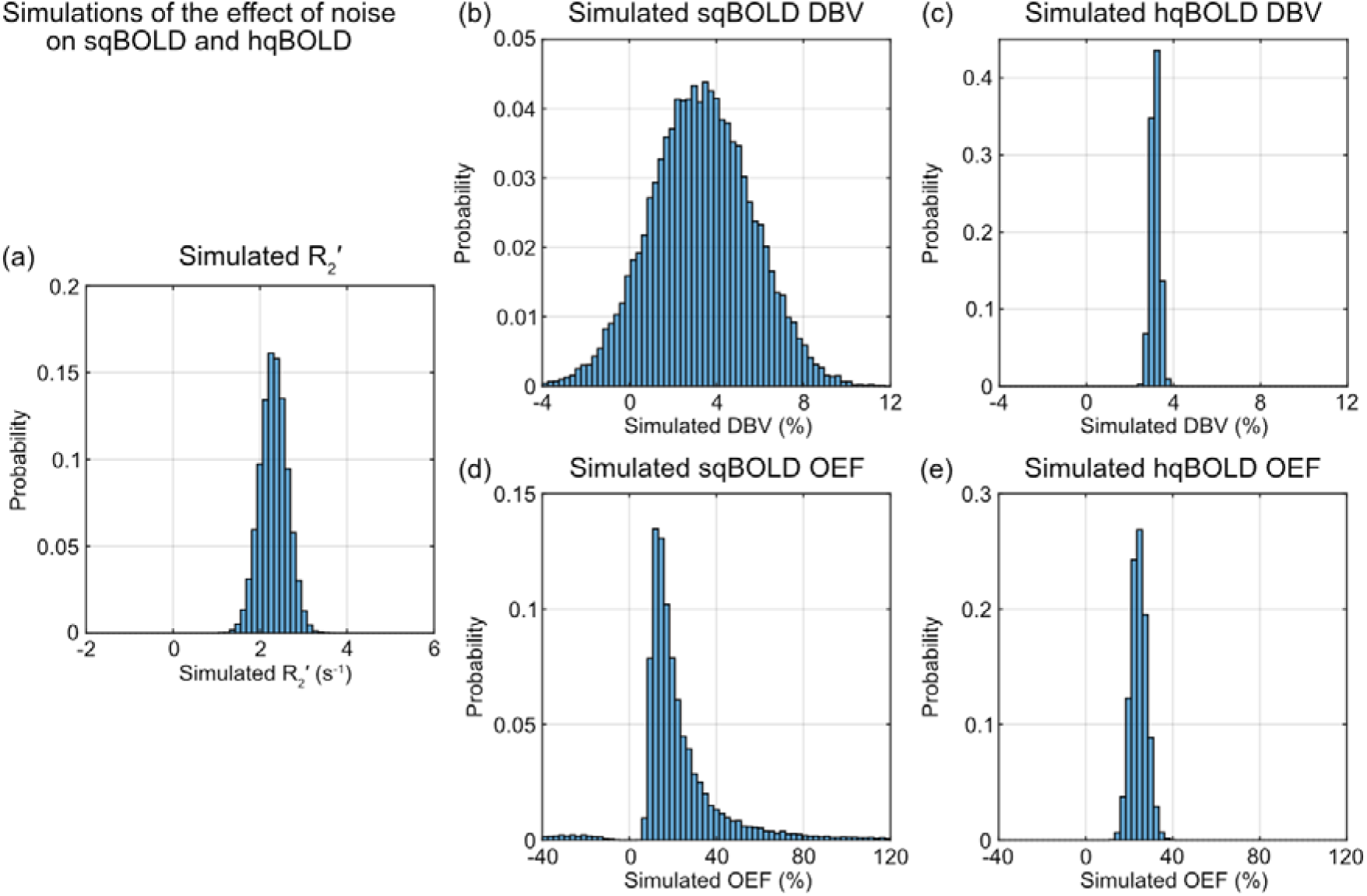
Simulations of the effect of noise for the two qBOLD approaches. (a) the distribution of R_2_′ values estimated from simulated qBOLD signals, (b-c) the distribution of deoxygenated blood volume (DBV) values for streamlined-qBOLD (sqBOLD) and hyperoxia-qBOLD (hqBOLD) techniques, respectively, and (d-e) the distribution of oxygen extraction fraction (OEF) values for sqBOLD and hqBOLD, respectively.

Figure 4 shows parameter maps of DBV and OEF from sqBOLD and hqBOLD for three slices (slices 2, 5 and 8) in a single subject (sub-02). A single slice from each of the ten participants is provided in Figure S1. DBV_hqBOLD_ maps demonstrate an obvious contrast between GM and WM, with lower values in WM as expected. In the GM the OEF_hqBOLD_ maps show physiologically plausible values for healthy brain tissue. However, in WM unphysiological values of OEF are produced with a mean OEF value of 140.9 %. In comparison, the DBV_sqBOLD_ and OEF_sqBOLD_ maps demonstrate little contrast between GM and WM and agree well with the initial implementation of this method^3^.

**Figure 4:**
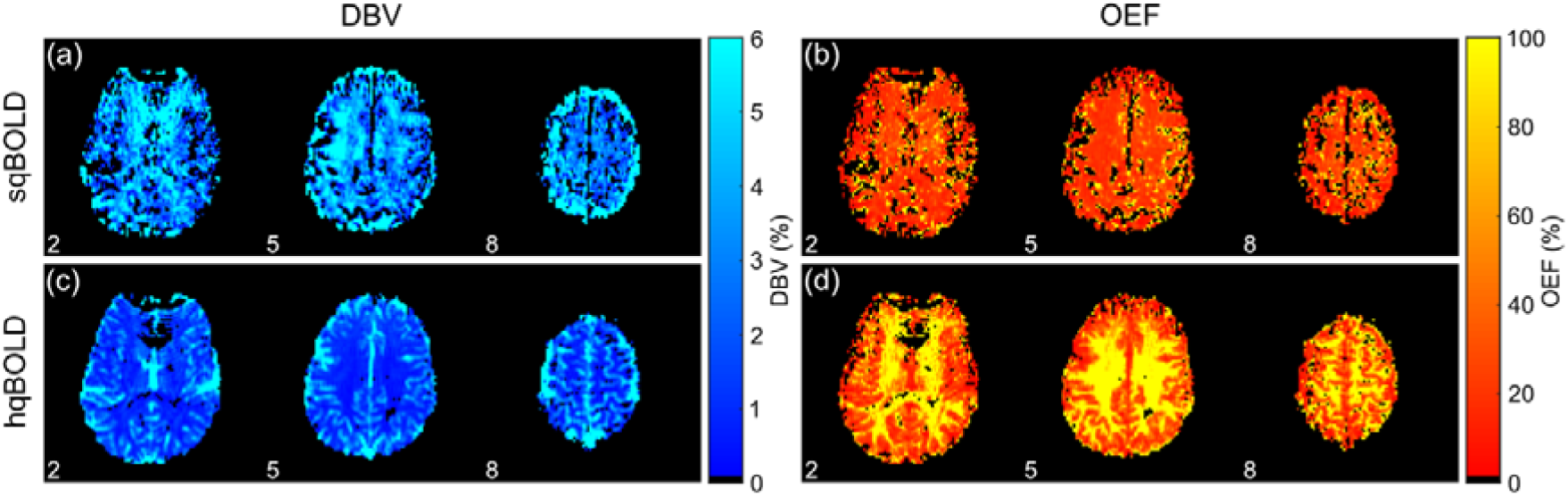
Example maps of deoxygenated blood volume (DBV) and oxygen extraction fraction (OEF) from the (a-b) the streamlined-qBOLD (sqBOLD) technique and (c-d) the hyperoxia-qBOLD (hqBOLD) method.

Table 3 shows regional estimates in GM extracted from hqBOLD and sqBOLD parameter maps. These estimates are compared to values of global OEF measured using TRUST. Group mean DBV_hqBOLD_ (1.8 ± 0.6 %) was found to be significantly different (p<0.05) to DBV_sqBOLD_ (3.9 ± 1.3 %). The one-way ANOVA was found to be significant (p<0.001) indicating that the mean OEF values for each group are not equal. Post-hoc pairwise comparisons revealed that OEF_sqBOLD_ is significantly different to OEF_hqBOLD_ (p < 0.001) and OEF_TRUST_ (p < 0.001), but that there was no significant difference between OEF_hqBOLD_ and OEF_TRUST_ (p = 0.77).

Figure 5 shows histograms of DBV_hqBOLD_, OEF_hqBOLD_, DBV_sqBOLD_ and OEF_sqBOLD_ voxel values within GM for one example participant (sub-03). These histograms show the distribution of voxel values that underlie the measurements presented in Table 3. DBV_hqBOLD_ exhibits a narrower distribution of voxel values compared to DBV_sqBOLD_ and OEF_hqBOLD_ displays a higher median OEF (Table 3) and a broader distribution of values compared to OEF_sqBOLD_. Histograms for all ten participants are presented in Figure S2.

**Figure 5:**
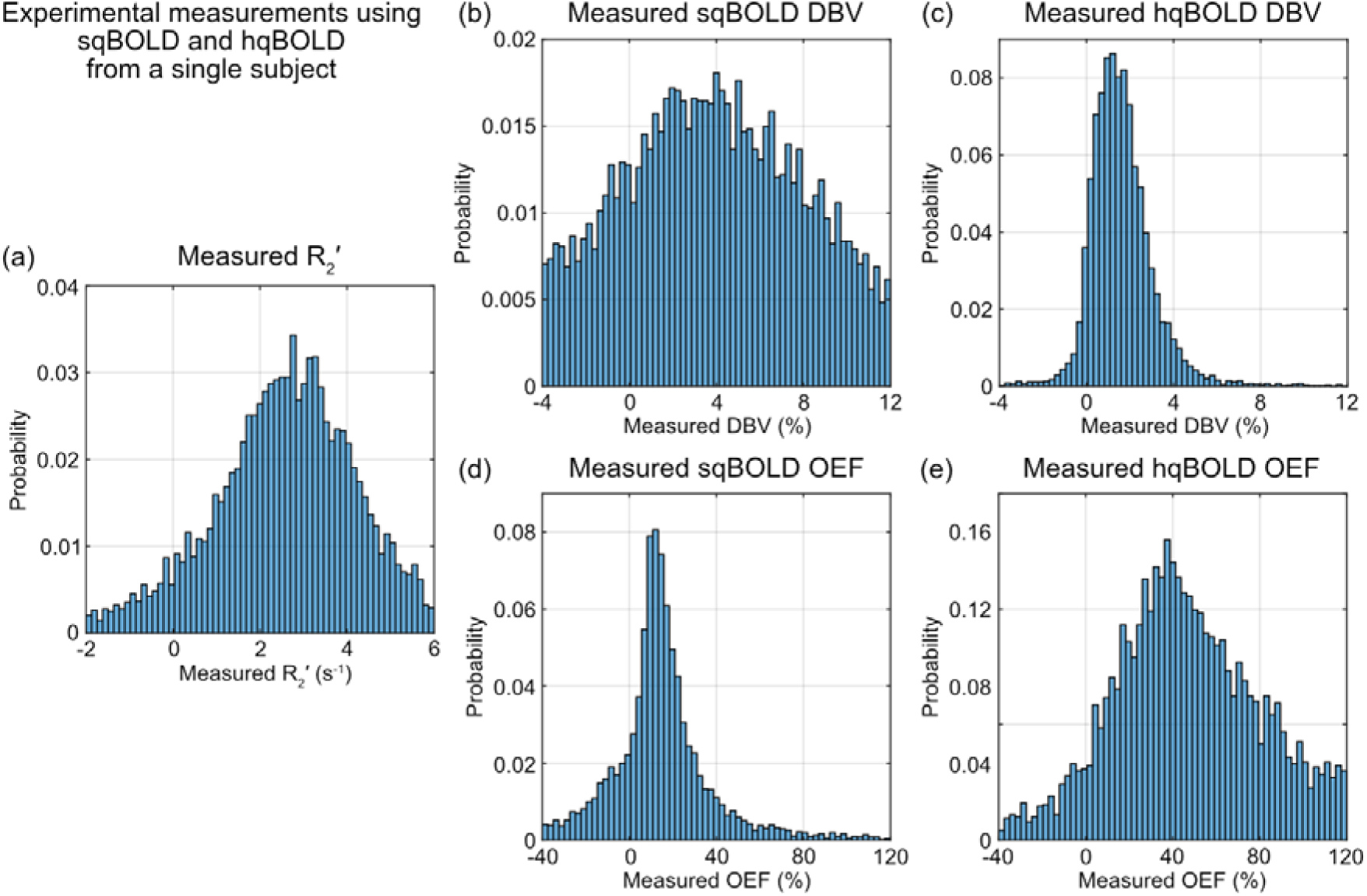
Histograms of experimental qBOLD measurements in grey matter. (a) the distribution of R_2_′ values, (b-c) the distribution of deoxygenated blood volume (DBV) values for streamlined-qBOLD (sqBOLD) and hyperoxia-qBOLD (hqBOLD) techniques, respectively, and (d-e) the distribution of oxygen extraction fraction (OEF) values for sqBOLD and hqBOLD, respectively.

Figure 6 shows OEF_TRUST_ compared to OEF_hqBOLD_ and OEF_sqBOLD_. Bland-Altman plots demonstrate a large bias between OEF_TRUST_ and OEF_sqBOLD_ of -24.3 percentage points whereas OEF_TRUST_ and OEF_hqBOLD_ are in better agreement with a smaller bias of -2.4 points between techniques.

**Figure 6:**
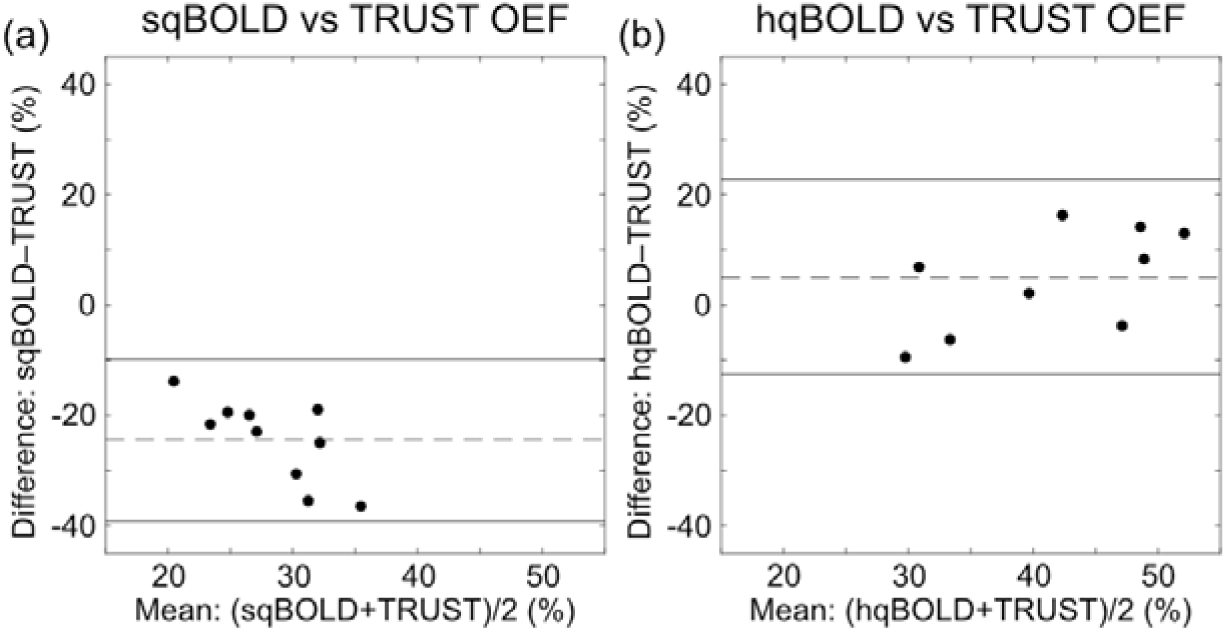
Bland-Altman plot comparing hyperoxia-qBOLD (hqBOLD) measures of oxygen extraction fraction (OEF) in global grey matter with streamlined-qBOLD (sqBOLD) and TRUST (T_2_ relaxation under spin tagging) measures of OEF.

In order to understand the unphysiological OEF in WM, Table 4 compares the ratio of the GM to WM values of R_2_′, DBV_sqBOLD_ and DBV_hqBOLD_. GM R_2_′ was found to be 22 % lower than WM, GM DBV_sqBOLD_ was found to be 29 % lower than WM and GM DBV_hqBOLD_ was found to be 234 % higher than WM.

## Discussion

The aim of this study was to investigate whether an independent measure of DBV based on the hyperoxia-BOLD signal could improve the accuracy of qBOLD measurements of OEF. Simulations of this new technique, referred to here as hqBOLD, predicted a considerable improvement in estimates of both DBV and OEF compared with the existing sqBOLD technique. Experimentally, measurements of OEF acquired using hqBOLD and sqBOLD were compared with whole brain oximetry measurements from TRUST. The hqBOLD measurements of OEF were found to be in the range 23 to 53%, in good agreement with literature values of OEF in healthy cortical GM (∼ 35 – 55 %)^40^. Whilst sqBOLD measurements of OEF were found to be systematically low (range 12 – 22 %) (Table 3). Furthermore, no significant difference was found between GM hqBOLD OEF measurements and TRUST OEF measurements (range 28 – 53 %).

### Simulations

An accelerated Monte Carlo framework which scales and combines results from single vessel radius simulations was used to examine systematic and noise related errors. Simulations of the sqBOLD technique were consistent with previous results^8^, namely that R_2_′ is linearly related to the true R_2_′ estimated from Eq. 3, that DBV_sqBOLD_ is a function of the OEF and this leads to a reduced dynamic range of OEF_sqBOLD_. One difference between this previous study and the current work is the definition of true DBV. As noted above, there isn’t a strict definition of DBV because the level of deoxygenation varies across the vasculature and the way that deoxygenation is translated into the BOLD signal is affected by the vessel radius. Hence DBV can be pragmatically described as either all vessels containing deoxyhaemoglobin (capillaries + veins) or only venous vessels. The impact of moving from the former definition in the previous study to the latter in the current study is a greater correspondence between the simulated measurement of R_2_′ with the true R_2_′ (slope 0.99). This would suggest that the deoxygenated blood in the capillaries only makes a small contribution to the measured R_2_′.

Simulations of DBV_hqBOLD_ provide an independent confirmation of the heuristic model (Eq. 8) for scaling the hyperoxia BOLD signal change to DBV. This heuristic was developed using an entirely different model to the one presented here based on analytical models of three vascular compartments^7^. In this previous work a narrower range of healthy OEF values were simulated (35 – 55 %). Despite this, Eq. 8 generalises well to the much larger OEF range simulated in this study (0 – 100 %), with minimal error in DBV_hqBOLD_ when OEF is greater than approximately 30 % (Fig. 2d). This is emphasised by the close correspondence between DBV_hqBOLD_ and true DBV with a slope of 0.95 for OEF > 30 % (Fig. 2c). Therefore, when combined with the simulated measurement of R_2_′ and Eq. 7, the dynamic range of OEF_hqBOLD_ is markedly improved.

Using noise levels estimated from the data, the simulations show that for a single physiological state the uncertainty in DBV_sqBOLD_ (Fig. 3b) is considerably larger than for DBV_hqBOLD_ (Fig. 3c). This might also explain the long tails and outliers for OEF_sqBOLD_ (Fig 3d and large excess kurtosis) as the reciprocal of the DBV distribution forms part of the OEF measurement (Eq. 9). When the shift of the distribution is small, or the uncertainty is large, the reciprocal of the distribution is bimodal and skewed. Hence negative values of OEF_sqBOLD_ are observed and skewed for positive values, unlike OEF_hqBOLD_ where the uncertainty in DBV is lower.

### Grey matter

Cortical GM measurements of DBV_hqBOLD_ (1.8±0.6 %) were significantly different (p<0.05) to DBV_sqBOLD_ derived measurements (3.9±1.3 %), with DBV_sqBOLD_ 2.2 times higher than DBV_hqBOLD_ on average. The sqBOLD measurements are consistent with previous implementations^3^ (3.6±0.4 %), whilst the hqBOLD estimates are slightly lower than past measurements^7^ (2.2±0.4 %). The hyperoxia-BOLD implementation in the current study differs in two ways from previous work: a shorter TR was used (down from 3 s to 1 s) and the hyperoxic gas mixture was presented differently. Automated gas delivery was used in this study, which benefits from isocapnic control and faster transitions between oxygen levels, compared with a two tube nasal cannula used in the previous study. An uncontrolled hyperoxic challenge, as in the latter case, can result in hyperventilation and associated hypocapnia^41^. However, hypocapnia should reduce the amplitude of the BOLD response to hyperoxia and hence we would expect the previous measurements to be lower than the current. Alternatively, a short TR could result in a decrease in the BOLD response to hyperoxia due to decrease in arterial blood T_1_ due to the presence of paramagnetic oxygen dissolved in arterial blood plasma. However, detailed study of this issue has predicted that T_1_ effects are negligible when compared with the changes in deoxyhaemoglobin elicited by hyperoxia^42^.

### Validation

The TRUST technique provides a way to measure whole brain OEF with few assumptions providing a way to benchmark sqBOLD and hqBOLD. Bland-Altman plots demonstrate that measurements of GM OEF made with hqBOLD agree well with TRUST oximetry across the group (Fig. 6b). The bias between the two methods was measured as -2.4 %. In contrast, sqBOLD measures of OEF in GM demonstrate poor agreement and a large bias of -24.3 % when compared to TRUST oximetry (Fig. 6a).

### White matter

The ratio of GM to WM DBV was measured as 0.7 for sqBOLD and 3.3 for hqBOLD. There are few studies in the literature that measure DBV quantitatively. A GM to WM DBV ratio of 2.29 was measured using qBOLD with the GESSE pulse sequence^2^. Alternatively, a study of CBVt measured a GM to WM CBVt ratio of 2.38. Both values are of a similar order to that measured using hyperoxia-BOLD in the current study. In contrast, DBV measured using sqBOLD in WM appears unphysiological given that it would suggest that WM has a higher vascular density than GM.

Under the assumptions: (i) that deoxyhaemoglobin is the dominant source of magnetic susceptibility in the voxel and (ii) that the blood vessels are uniformly and randomly distributed, Eq. (3) describes R_2_′ as a function of OEF and DBV. Furthermore, OEF is generally found to be the same in GM and WM in PET measurements^43^. Therefore, under these conditions the ratio of GM to WM R_2_′ would be expected to scale in the same way as DBV. Hence, we would expect the R_2_′ of GM to be 2.29 – 3.31 times higher than for WM, but the ratio was measured to be 0.78. A recent investigation of R_2_′ in WM has found that the two assumptions above are likely to be violated^44^. This study followed a multiparametric qBOLD approach whereby R_2_′ is estimated from maps of R_2_ and R_2_* and estimated the angle between the primary eigenvector of WM voxels and B_0_ using DTI. R_2_′ was found to increase as the angle increased. This orientation effect was considered by the authors to be a function of the preferential alignment of blood vessels parallel to WM fibre tracks and the magnetic susceptibility effects arising from the myelin rich WM fibres. Whilst orientation undoubtedly has a role at the voxel level, in the measurements presented in the current study this is likely to have been averaged out over the WM ROI. However, the additional magnetic susceptibility effect from myelin will remain and contribute to the measured R_2_′.

### Limitations and future work

It is worth noting some of the relevant limitations of the protocols used in this study. Firstly, the discrepancy in total acquisition time between sqBOLD (9:12 mins) and hqBOLD (29:12 mins) may at first glance appear to be an unfair comparison from an SNR perspective.

However, as the simulations show, the underestimation of OEF is not due to a limitation in SNR but is caused by a systematic error. This study focussed on reducing this systematic error and future efforts will be directed towards reducing the overall scan duration.

The sqBOLD protocol used here has only minor differences compared with previous studies i.e. a slightly higher density of sampling at long τ values. We have shown that the long TE used here (TE = 80 ms) results in increased signal loss at the spin echo, due to the scale of the vessel distribution approaching the diffusion narrowing regime, causing DBV to be overestimated^8^. Unfortunately acquisition of the data in this study commenced before this simulation study had been completed and hence any improvements could not be implemented. Similarly, we have developed a Bayesian analysis tool for sqBOLD data, which has been shown to reduce the variance in parameter estimates^5^. However, since a tool to analyse the hqBOLD data in a Bayesian framework was not available, we chose to use least squares fitting for both techniques. However, we remain hopeful that an optimised sqBOLD protocol coupled with Bayesian model fitting can overcome some of the inaccuracy of the protocol used here.

The TRUST protocol used in this study had a longer echo time (7 ms) than recommended in a recent study^45^ (3.6 ms). This has been shown to result in an overestimation of T_2b_ that leads to a slight overestimation in OEF of approximately 3-4%. However, this overestimation was shown to be dependent on image SNR and hence the number of averages acquired in this study was increased from the recommended number of three to four.

## Conclusion

It has been shown by simulation and experiment that introducing an independent measure of DBV into the qBOLD framework provides improved estimates of OEF. Whilst mq-BOLD has previously used an independent measure of CBVt from DSC, this is not the relevant blood volume in the context of qBOLD. Hyperoxia-BOLD provides a quantitative and specific measure of DBV, such that the quantification of OEF is also improved. With further refinement this method has the potential to provide a clinically relevant measure of tissue oxygenation.

## Appendix A

The code used to generate the simulation results can be downloaded from the Zenodo repository, doi: https://doi.org/10.5281/zenodo.14812818. The raw imaging data can be accessed via the Oxford Research Archive repository, doi: http://doi.org/10.5287/bodleian:X59eGb0pv. The code used to analyse these can be accessed via the Zenodo archive, doi: https://doi.org/10.5281/zenodo.14814607.

## Appendix B: Supplementary data

Supplementary data to this article can be found online at <INSERT DOI TO SUPPLEMENTARY DATA>.

## Acknowledgements

This work was supported by the Engineering and Physical Sciences Research Council [grant number EP/K025716/1].

---

**Figure S1:**
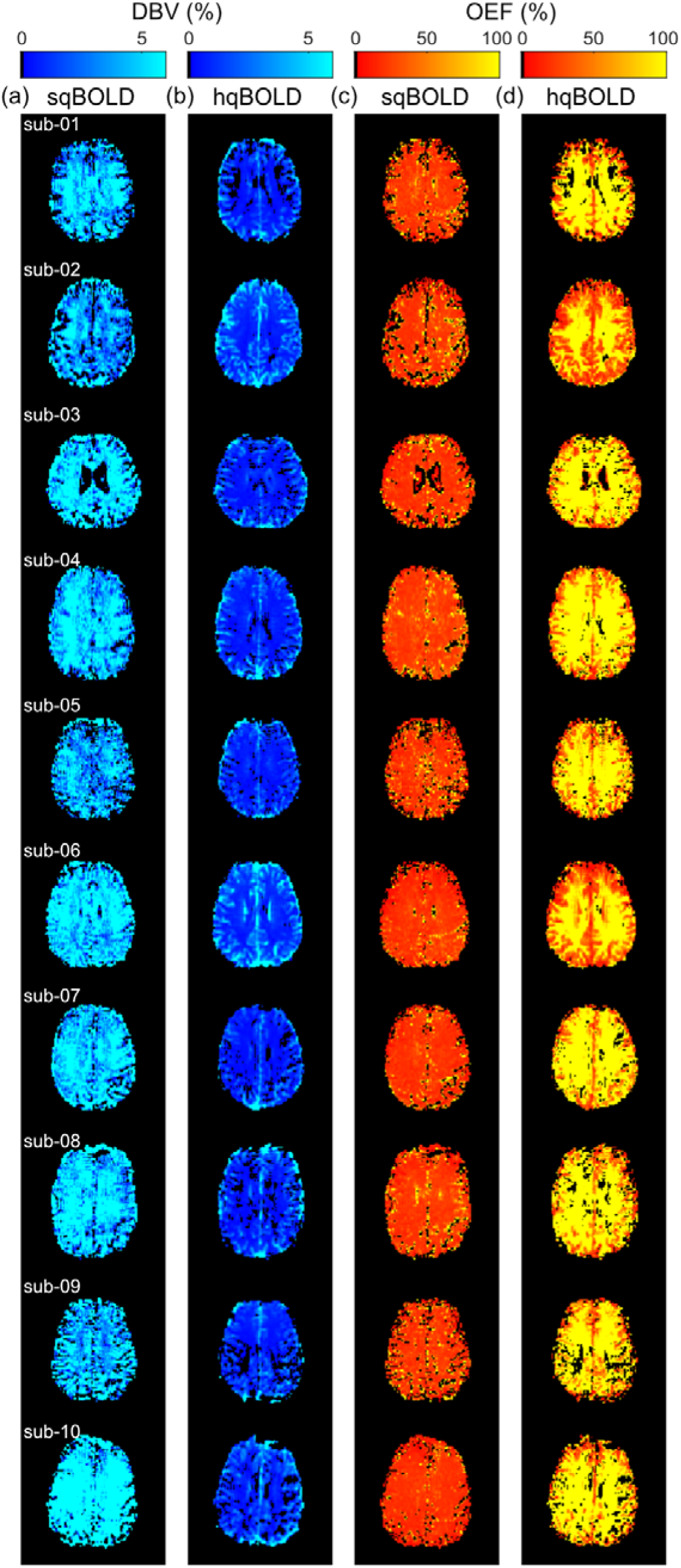
A single slice of the streamlined-qBOLD (sqBOLD) and hyperoxia-qBOLD (hqBOLD) data of each of the participants. (a-b) deoxygenated blood volume (DBV) and (c-d) oxygen extraction fraction (OEF) maps.

**Figure S2:**
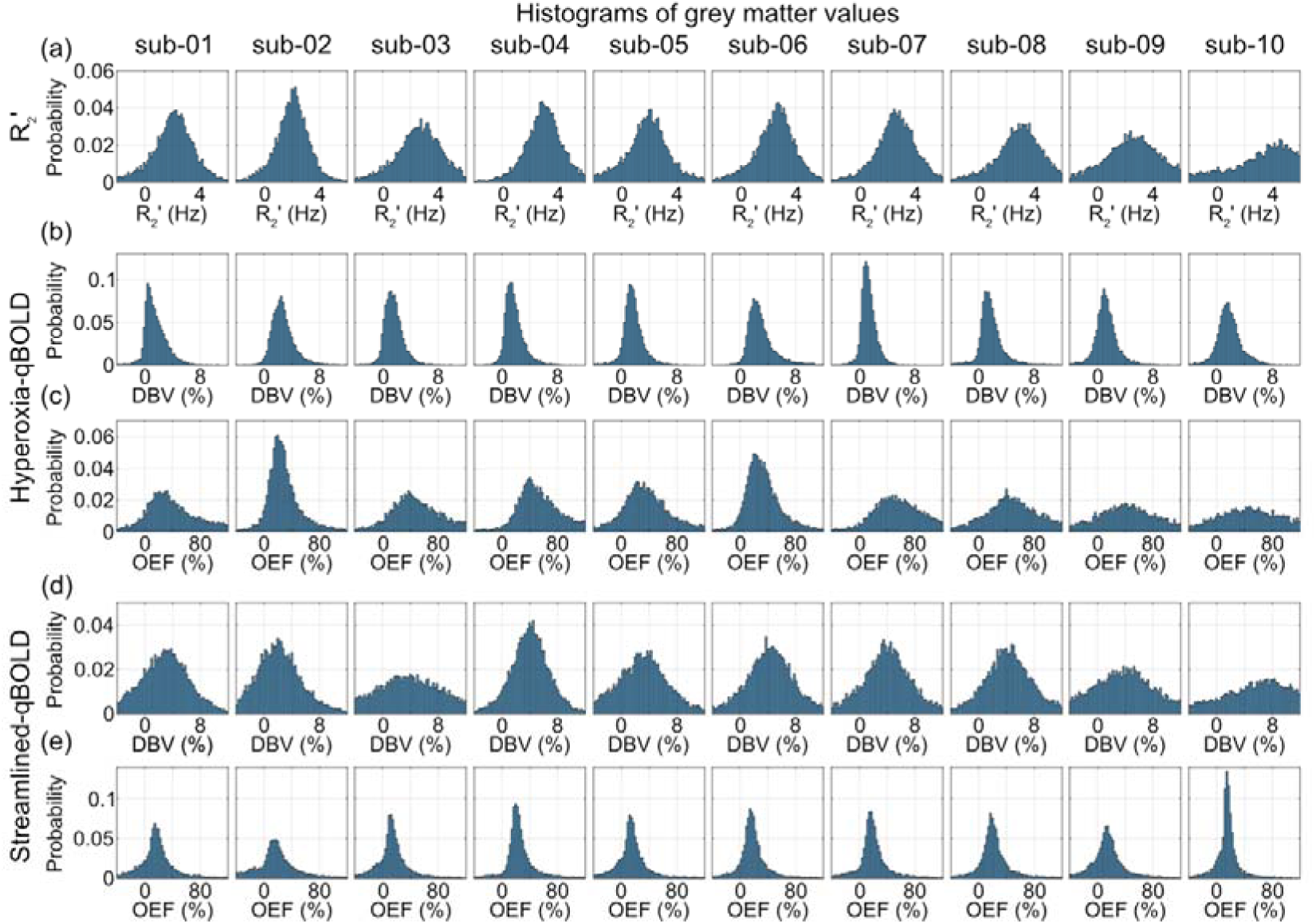
Histograms of experimental qBOLD measurements in grey matter for all subjects. (a) the distribution of R_2_′ values, (b-c) the distribution of deoxygenated blood volume (DBV) and oxygen extraction fraction (OEF) values for hyperoxia-qBOLD (hqBOLD) and (d-e) the distribution of DBV and OEF values for streamlined-qBOLD (sqBOLD).

## Notes

https://doi.org/10.5281/zenodo.14812818

http://doi.org/10.5287/bodleian:X59eGb0pv

https://doi.org/10.5281/zenodo.14814607

### Competing Interest Statement

The authors have declared no competing interest.

### Summary of Updates

This revision of the manuscript refocusses the paper to concentrate on grey matter estimates of oxygen extraction fraction and supplements these experimental measurements with simulations comparing streamlined-qBOLD with the newly developed hyperopia-qBOLD technique.

