## Supplemental data for "Improving qBOLD based measures of oxygen extraction fraction using hyperoxia-BOLD derived measures of blood volume"

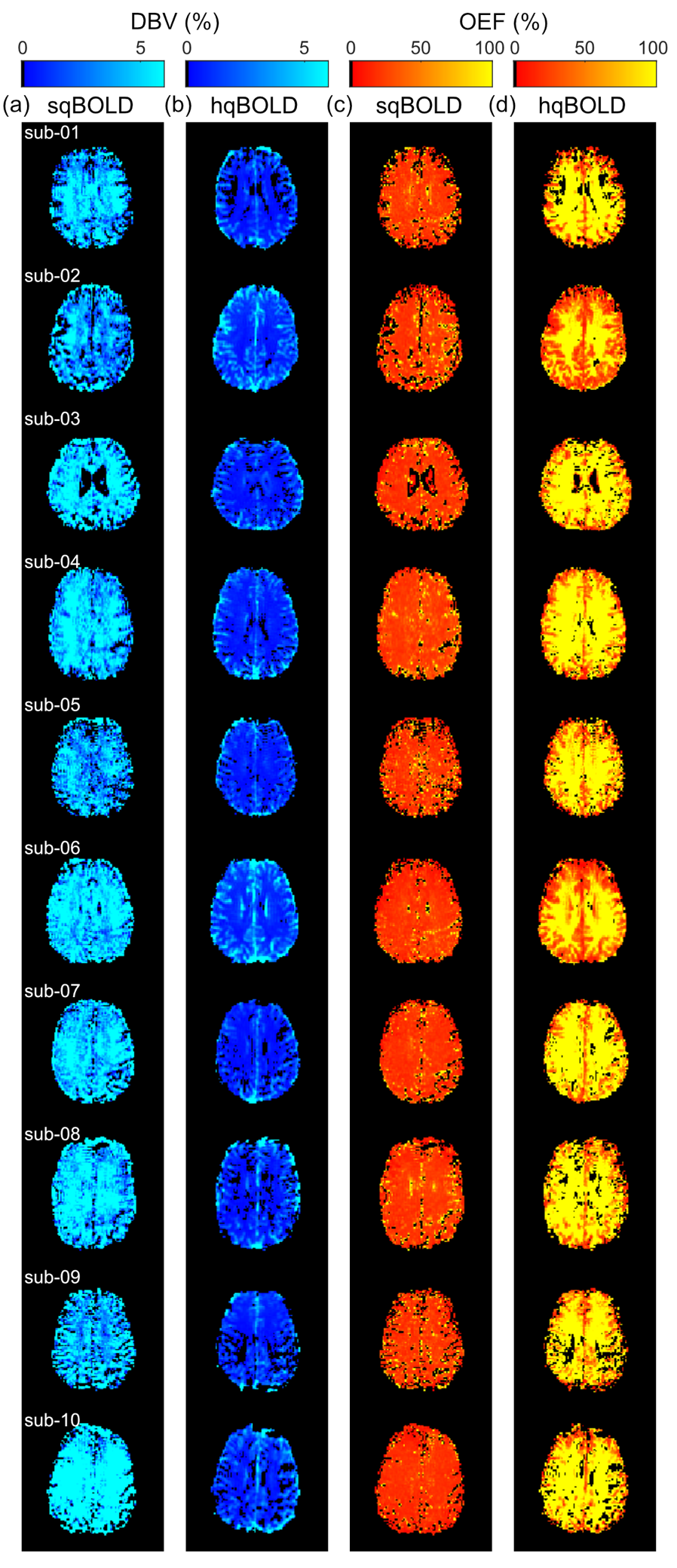

**FIGURE S1** A single slice of the streamlined-qBOLD (sqBOLD) and hyperoxia-qBOLD (hqBOLD) data of each of the participants. (a-b) deoxygenated blood volume (DBV) and (c-d) oxygen extraction fraction (OEF) maps.

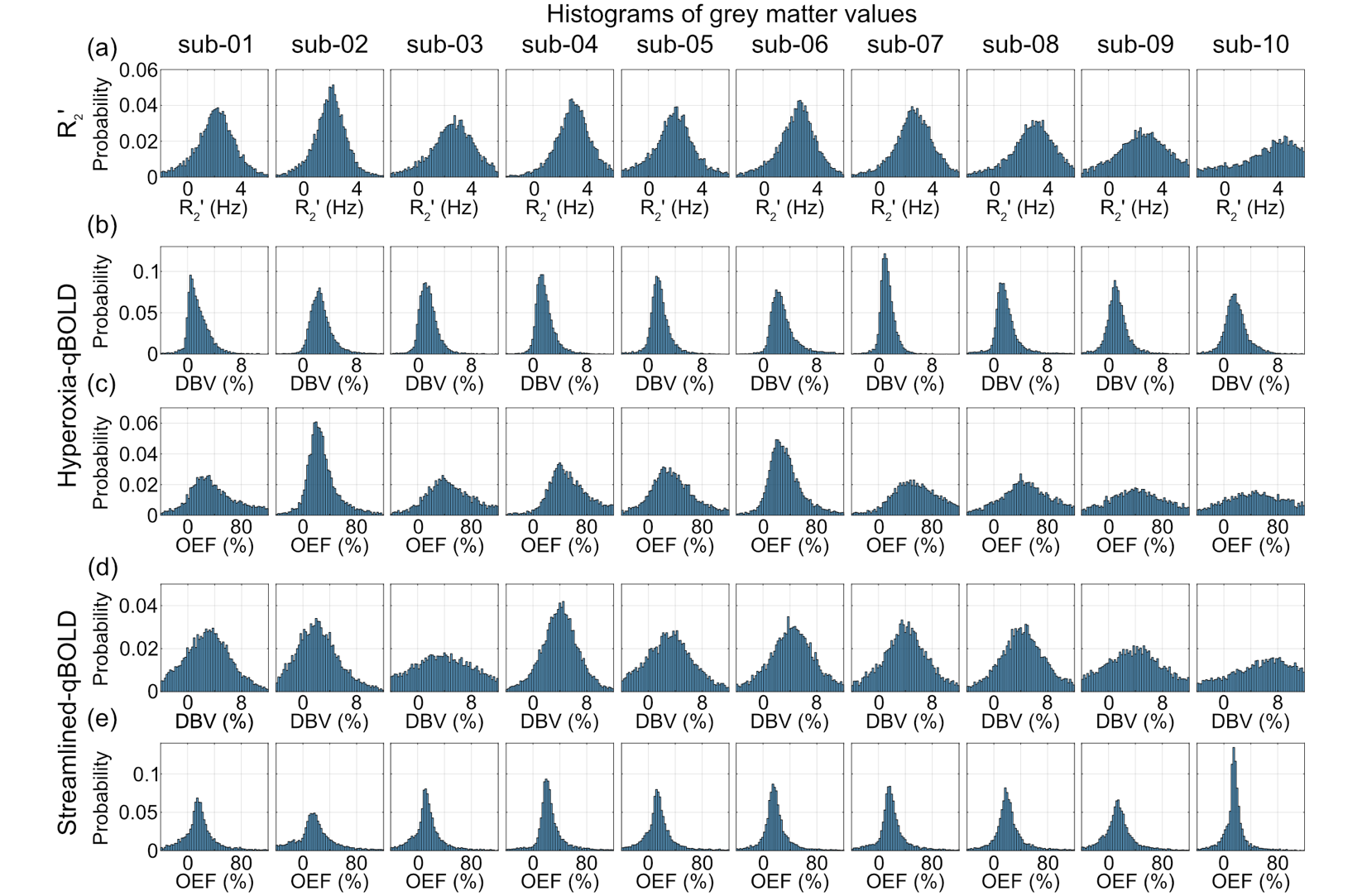

**FIGURE S2** Histograms of experimental qBOLD measurements in grey matter for all subjects. (a) the distribution of R_2_′ values, (b-c) the distribution of deoxygenated blood volume (DBV) and oxygen extraction fraction (OEF) values for hyperoxia-qBOLD (hqBOLD) and (d-e) the distribution of DBV and OEF values for streamlined-qBOLD (sqBOLD).
